# Neutral Sphingomyelinase-2 Restrains TAZ to Suppress Breast Tumor Growth

**DOI:** 10.64898/2026.05.13.724348

**Authors:** Andrew E. Resnick, Victoria Franzi, Botheina K. Ghandour, Sam B. Chiappone, Sebastien Lalanne, Monica E. Alexander, Deanna M. Peperno, Jowana Obeid, Ishmam Pritam, Ian D. Miranda, Nicolas Coant, Michael V. Airola, Mehdi Damaghi, Fabiola N. Velazquez, Yusuf A. Hannun, Christopher J. Clarke

## Abstract

Loss of tumor suppressor gene (TSG) activity is pervasive across cancers and linked to worse clinical outcomes, yet therapeutic efforts aimed at restoring TSGs have remained elusive. One underexplored avenue to address this problem is the targeting of metabolic signaling pathways that actively enforce tumor suppressive programs. Ceramide (Cer), the central hub of the sphingolipid (SL) metabolic network, has long been thought to have tumor suppressive functions, though its mechanistic roles remain incompletely defined. Here, we identify neutral sphingomyelinase-2 (nSMase2) as a critical mediator of Cer-dependent tumor suppression. We show that nSMase2 is frequently suppressed in breast cancer (BC) and its restoration inhibits tumorigenesis. Biologically, this was linked to the suppression of anchorage-independent growth (AIG) and to restraint of the HIPPO pathway effector TAZ, but not its paralog YAP. Taken together, these findings define a previously unrecognized metabolic tumor suppressor pathway, clarify ambiguities in both SL and HIPPO signaling networks, and highlight reactivation of nSMase2-Cer signaling as a potential therapeutic strategy in BC.

---

Cancer is a complex, multistep disease driven by progressive metabolic reprogramming (1) that fundamentally reflects an imbalance in cellular signaling/regulatory networks through both the activation of oncogenic drivers (2) and the inactivation of tumor suppressors (2). Loss of tumor suppressor gene (TSG) activity is pervasive across cancers (3) and arises through diverse mechanisms including copy number loss (RB1) (4), mutation (TP53) (5), and epigenetic repression (HIC1) (6). And unlike oncogenic drivers, disruption of TSG signaling typically requires loss of both functional alleles (7). Biologically, loss of TSG function has been linked with aggressive phenotypes including therapeutic resistance (8), genomic instability (9, 10), and immune evasion (11). Consequently, TSG alterations are often clinically associated with worse outcomes (12, 13). While restoring the signaling of functionally inactivated TSGs has been considered as a promising approach for treating aggressive cancers (14, 15), therapeutic reactivation of TSG pathways has proven challenging compared to targeting oncogenic signaling (14, 16), largely due to issues of specificity (17), context dependence (18), and druggability (17). These limitations highlight the need to identify tumor suppressive signaling networks that are both mechanistically defined and therapeutically tractable. Important underexplored opportunities lie in metabolic signaling pathways that actively enforce tumor suppressive programs. One such emerging pathway is that of sphingolipid (SL) metabolism.

Sphingolipids (SLs) are a family of lipids implicated in many biological effects relevant to tumorigenesis (19), and the dysregulation of SL metabolism has been observed across multiple cancers (20–23). Ceramide (Cer), the central hub of the SL metabolic network (19) has long been thought to function as a tumor suppressor lipid (24) due to its anti-proliferative (25, 26) and pro-apoptotic (24, 27) properties. However, intracellular Cer is generated by multiple biochemical pathways (19) including *de novo* SL synthesis, sphingomyelin hydrolysis by sphingomyelinases (SMases), and salvage-pathway based recycling of sphingosine (Sph) from turnover of complex SLs. More critically, each of these pathways generates Cer pools with divergent biological effects (19, 28). While the inhibition of *de novo* synthesis of Cer has been shown to promote autophagy (29), the hydrolysis of sphingomyelin into Cer has been implicated in anti-tumor biologies such as contact inhibition (30), growth arrest (30, 31), and programmed cell death (32–35). Therefore, to harness Cer therapeutically, it is essential to define the specific pathways and relevant enzymes of SL metabolism involved.

Neutral SMase-2 (nSMase2), a key regulated enzyme that hydrolyzes sphingomyelin into Cer, is the best studied of the four known mammalian neutral SMase isoforms (36–41). We and others have implicated nSMase2 in anti-tumor biologies (30–35) and demonstrated regulation by anti-cancer agents such as all-trans retinoic acid (31) and anthracyclines (42). Early evidence suggesting alterations of nSMase2 in cancer came from localization of the gene encoding nSMase2, *SMPD3*, on chromosomal locus 16q22.1 (43). This is a frequent site of loss of heterozygosity across multiple malignancies (44–46), including breast cancer (BC) (47), and correlated strongly with advanced disease and worse clinical outcomes (44, 47). Subsequent studies discovered that genetic loss of nSMase2 activity in mice lead to spontaneous liver tumor formation (48) and gain of nSMase2 function is associated with an increased response to PD-1 immunotherapy in melanoma (49). The former study in liver cancer was the first to suggest *SMPD3* as a TSG, the latter study in melanoma adding to the anti-cancer role of *SMPD3*. Beyond genomic loss, the *SMPD3* gene is subject to epigenetic silencing in several aggressive cancers including BC (31, 50), hepatocellular carcinoma (51), lung cancer (52), renal cancer (53), oral carcinoma (54). However, while decreased nSMase2 expression has been observed in recurrent HER2-positive BC tumors relative to matched primary lesions (55), studies in basal-like BC have reported that nSMase2 can promote metastatic phenotypes by inducing the packing of pro-angiogenic microRNAs into exosomes (56). Similarly, targeting exosome release with the nSMase2 inhibitor GW4869 in HER2-positive BC cells led to an increased response to trastuzumab therapy. Collectively, these findings point to a multifaceted and incompletely understood role for nSMase2 in BC biology. More critically, the molecular mechanisms that determine when and how nSMase2 constrains tumor growth and which downstream signaling pathways mediate these effects remain undefined. Resolving this uncertainty is essential, given the central position of Cer within SL metabolism and the diverse cellular outcomes associated with altered SL signaling.

In this work, we demonstrate that nSMase2 is frequently suppressed in basal-like and HER2-positive tumors and that restoration of nSMase2 activity functions as a potent tumor suppressor in these aggressive cancers. Using both *in vitro* and *in vivo* model systems integrated with transcriptomic profiling, we demonstrate that nSMase2 selectively suppresses anchorage-independent growth (AIG), a hallmark of malignant transformation and tumorigenicity. Mechanistically, the nSMase2-Cer axis activates the HIPPO pathway kinase LATS, leading to the phosphorylation and subsequent cytoplasmic sequestration of TAZ, but not YAP. Together, these findings define a previously unrecognized nSMase2–Cer–LATS–TAZ signaling axis, resolve long-standing gaps in both context-specific Cer function as well as resolving YAP vs TAZ dependent signaling programs, and positions nSMase2-Cer signaling as a tractable vulnerability in aggressive BCs.

## MATERIALS AND METHODS

### Reagents

Standard laboratory reagents were from Life Technologies, BioRad, Millipore Sigma or Fisher Scientific unless otherwise specified. Key reagents included: Lipofectamine 2000 (#11668019, Life Technologies), polybrene (#TR-1003-G, Millipore Sigma), puromycin and blasticidin (InvivoGen, San Diego, CA). Complete antibody information, including all primary antibodies, dilutions, and catalog numbers, is provided in the Key Resources Table.

### Cell lines and cell culture

MDA-MB-231 (ATCC HTB-26) and JIMT-1 (DSMZ ACC-589) cells were obtained from the American Type Culture Collection (Manassas, VA, USA) and DSMZ-German Collection of Microorganisms and Cell Cultures GmbH (Braunschweig, Germany), respectively, and authenticated by STR profiling. Cells were cultured in Dulbecco’s Modified Eagle’s Medium (DMEM; #11965092, Gibco/Life Technologies) supplemented with 10% (v/v) heat-inactivated fetal bovine serum (FBS; #26400044, Life Technologies). Cell lines were maintained at 37°C, 5% CO₂, and passaged at ∼85% confluence using Trypsin-EDTA (0.05%; #25300054, Gibco/Life Technologies) following washing with Dulbecco’s Phosphate-Buffered Saline (DPBS; #14190250, Life Technologies). Cells were confirmed to be mycoplasma-free monthly using a luminescence-based assay. STR profiling was performed every 6 months to ensure cell line identity. All experiments were conducted using cells between passages 5 and 25.

### Plasmids

*p*CMV-VSV-G (Addgene #8454) and pCMV-dR8.2 DVPR (Addgene #8455) were gifts from Robert Weinberg (MIT). pLenti CMV/TO Puro (Addgene #17293) was a gift from Eric Campeau (MISO Chip). Custom pLenti CMV vectors containing SMPD3-WT, SMPD3-H639A (a catalytically inactive mutant), YAP1-5SA, and WWTR1-4SA (constitutively active mutants) were generated via TWIST biosciences and Genscript. All plasmids were verified by Sanger sequencing prior to use.

### Generation of stable cell lines

To generate lentiviral particles, 2×10⁶ HEK293T cells were seeded in 100 mm dishes. After 24 h, cells were co-transfected with 1.5 µg of the target plasmid, 1.5 µg of pCMV-VSV-G, and 1.5 µg of pCMV-dR8.2 DVPR using 13.5 µL of Lipofectamine 2000 in 600 µL Opti-MEM (#31985062, Life Technologies) according to the manufacturer’s protocol. After overnight incubation, the medium was changed to fresh DMEM with 10% FBS. At 72 h post-transfection, virus-containing media were harvested, filtered through a 0.45 µm PVDF syringe filter (#SLHVR33RS, Millipore Sigma), aliquoted, and stored at -80°C. Freeze-thaw cycles were avoided. For infection, MDA-MB-231 and JIMT-1 cells were seeded at 1×10⁵ cells per well in 6-well plates. The next day, cells were incubated with 1 mL viral supernatant in 1 mL of growth media containing polybrene (16 µg/mL). After 24 h, the medium was replaced with fresh growth medium. After another 24 h, cells were trypsinized and cultured in media containing blasticidin (10 µg/mL) or puromycin (1 µg/mL) for 7 days to select for stably transduced cells. Optimal antibiotic concentrations were predetermined for each cell line using kill curves. Polyclonal populations were used for all experiments, and stable overexpression was confirmed by immunoblot, immunofluorescence, and/or RT-qPCR.

### Analysis of gene expression by quantitative RT-PCR and microarray

For cells, total RNA was extracted using the PureLink RNA Mini Kit (#12183018A, Life Technologies) according to the manufacturer’s protocol. RNA concentration and purity were measured via Nanodrop spectrophotometry. 1 µg of total RNA was reverse transcribed to cDNA using the SuperScript III First-Strand Synthesis System (#11752250, Life Technologies). Real-time qPCR was performed on an ABI 7500 Fast system using iTaq universal probes supermix (#1725135, BioRad) and pre-designed TaqMan Gene Expression Assays (Thermo Fisher; see Key Resources Table for IDs). Relative gene expression was calculated using the ΔΔCt method, normalizing to the geometric mean of three reference genes: *ACTB*, *GAPDH*, and *18S*. All experiments were performed with three biological replicates, each with technical triplicates. For xenograft tumors, frozen samples were homogenized in TRIzol Reagent (Invitrogen, #15596026) using a tissue homogenizer (Qiagen TissueRuptor I, #305273585869). Total RNA was purified using the PureLink RNA Mini Kit with on-column DNase treatment (Qiagen, #79254). Samples were sent to the Stony Brook Genomics Core for RNA quality control and for Human Affymetrix GeneChip Clarium S microarray. Raw data was processed using the RMA algorithm in the Affymetrix Transcriptome Analysis Console. Gene-level summarization and batch correction (ComBat) were performed. Microarray data will be deposited in the Gene Expression Omnibus (GEO).

### Analysis of cellular protein by SDS-PAGE and Immunoblot

Cells were lysed in RIPA buffer supplemented with a protease inhibitor cocktail (#501657350, Millipore Sigma). Protein concentration was determined using the Bradford assay. Equal protein amounts (10-20 µg) were separated by SDS-PAGE on 4-20% gradient gels (#WXP42020BOXA, Life Technologies) using Criterion Cell Tanks (#1656001, Bio-Rad) and transferred to 0.45 μm nitrocellulose membranes (#1620115, Thermo Fisher) using a Criterion Blotter (#1704070, Bio-Rad). Membranes were blocked for 1 h in 5% non-fat dry milk in PBS with 0.1% Tween-20 (PBST), then incubated with primary antibodies overnight at 4°C. After washing, membranes were incubated with HRP-conjugated secondary antibodies for 1 h at room temperature. Protein bands were detected using Pierce ECL Western Blotting Substrate (#32106, Thermo Fisher). β-actin or GAPDH served as a loading control. All blots were repeated in at least three biological replicates.

### Neutral sphingomyelinase (nSMase) activity assay

N-SMase activity was assayed *in vitro* as described previously (30) utilizing ^14^C-[methyl]sphingomyelin as substrate.

### Analysis of cellular sphingolipids

For all lipid experiments, cells were washed, scraped in ethyl acetate/isopropyl alcohol/water (60:30:10, v/v/v), extracted via Bligh & Dyer Method, and submitted to the stony brook lipidomics core facility for tandem tandem LC/MS mass spectrometry. Lipid levels were normalized to the total lipid phosphate level of the sample as described previously (57).

### Cell viability and proliferation assays

For metabolic activity assays (MTT), 1×10^5^ (MDA-231) or 7.5×10^4^ (JIMT-1) cells were seeded in 6-well plates. After overnight attachment, cells were incubated with 1:1 Media:Thiazolyl Blue Tetrazolium Bromide (MTT; #97062-380, VWR International; 5 mg/mL in DPBS) for 30 min at 37°C. The medium was aspirated, and formazan crystals were dissolved in DMSO. Optical density was measured at 570 nm using a SpectraMax M5 plate reader (Molecular Devices). For direct viability assays, 2×10⁵ cells were seeded and harvested at the indicated time points. The cell suspension was mixed 1:1 with trypan blue solution (#15250061, Fisher Scientific), and viable and non-viable cells were counted using a hemocytometer. Data are expressed as the percentage of viable cells or as total viable cell number.

### Anchorage-independent survival and soft agar assays

To assess anchorage-independent survival, 2×10⁵ cells were cultured in ultra-low attachment 6-well plates (#07200601, Fisher Scientific) for 48 h. Suspension cells were harvested, dissociated with trypsin, and viability was assessed by trypan blue exclusion as described above. To assess anchorage-independent growth via soft agar colony formation, a base layer of 0.6% bacteriological agar (#97064-336, VWR International) in growth medium was prepared in 6-well plates. Cells (2×10⁵ MDA-MB-231 or 7.5×10⁴ JIMT-1) were suspended in 0.3% agar in growth medium and plated on top of the solidified base layer. Cultures were fed every 3-4 days with fresh medium and incubated for 2-4 weeks. At the endpoint, colonies were stained overnight with Nitro Blue Tetrazolium Chloride (#N6495, Fisher Scientific; 1 mg/mL). Plates were imaged and colonies were counted automatically using a Nexcelom Celigo Image Cytometer. A size threshold of ≥50 µm in diameter was used to define a colony.

### Analysis of nSMase2 levels by immunohistochemistry

The nSMase2 mouse monoclonal antibody was generated at the Medical University of South Carolina Antibody Facility using a highly purified (>95%) recombinant nSMase2 fragment (residues 113-655) as an antigen. Antibody was validated with JIMT EV and N2- expressing tumors. Tissue microarray (obtained from US Biomax; BR721, T086c) immunohistochemistry experiments were performed as described previously (58).

### 3D Matrigel culture assay

Growth factor-reduced Matrigel (80 µL; Corning) was plated in 4-well chamber slides and allowed to solidify (37°C, 15 min). Cells (4×10⁴ MDA-MB-231 or 6×10⁴ JIMT-1) were seeded onto the Matrigel and overlaid with 800 µL of DMEM containing 4% Matrigel. The medium was replaced every 4 days for 14 days. At the endpoint, spheroids were processed for western blot as described.

### 3D Spheroid Generation, Embedding, and Cryosectioning

To generate spheroids, (2×10^3^) cells were seeded in sterile, ultra-low-attachment U-bottom 96-well plates and incubated under standard culture conditions (37°C, 5% CO₂). One day after seeding, plates were centrifuged (20×g, 5 minutes, 25 °C) to promote cell aggregation, followed by an additional 2 days of incubation. The culture medium was then replaced and supplemented with 5% Matrigel (#356237, Corning) and plates were centrifuged again at 20 × g for 5 minutes at 25 °C. Six days after Matrigel addition, spheroids were treated with a LATS kinase inhibitor (5 µM) for 48 hours. Following treatment, spheroids were washed with PBS, fixed in 4% paraformaldehyde (PFA), and embedded in 15% gelatin using plastic molds. Samples were stored at −80 °C until further processing. Frozen blocks were mounted in carboxymethyl cellulose (CMC) and equilibrated in a CM1950 cryostat (Leica Biosystems) at −20 °C for 15 minutes prior to sectioning. Sequential 10 µm sections were collected for immunofluorescence analysis.

### Xenograft studies

Eight-week-old female NSG (NOD.Cg-Prkdcscid Il2rgtm1Wjl/SzJ) mice (The Jackson Laboratory, #005557) were used. Following a 7-day acclimation period, mice were randomized into experimental groups and orthotopically implanted with 2×10⁶ cells suspended in 50 µL of 1:1 DPBS:Matrigel into the 4th mammary fat pad. Tumor volume was measured twice weekly with digital calipers. Estimated ellipsoid tumor volume during monitoring was calculated as ((ν × Length × Width²)/6); final tumor volumes were calculated as ((4π × Length × Width × Height)/18) to account for 3-dimensional measurements. At endpoint (predefined as tumor volume >1000 mm³ or signs of distress), mice were euthanized. Primary tumors and lungs were harvested for analysis. Lungs were fixed in 10% formalin, paraffin-embedded, sectioned, and stained with H&E. Metastatic burden was quantified from digital images of lung sections (imaged on a Zeiss AXIO Imager M2) by measuring the percentage of total lung area occupied by tumor nodules using ImageJ. Quantification was performed on at least three sections per lung, spaced 100 µm apart. Sample sizes were calculated a priori to detect 30% differences in tumor volume with 95% power at α=0.05.

### Analysis of subcellular localization by immunofluorescence

Cells were fixed with 2% paraformaldehyde (10 min, 37°C), permeabilized with ice-cold 100% methanol (10 min, -20°C), washed with PBS, and blocked with 2% human serum in PBS for 1 h at room temperature. Tumor samples were processed as described (58). Primary antibodies (e.g., anti-FLAG, anti-V5, anti-YAP, anti-TAZ; 1:200) were diluted in blocking buffer and incubated (2 h, RT for cells; overnight, 4°C for tumors). After washing, cells were incubated with Alexa Fluor 488-conjugated secondary antibodies (1:200) for 1 h at 4°C in the dark. Nuclei were counterstained with DAPI. Secondary antibody-only controls were included to confirm signal specificity. Images were captured on an Olympus IX-70 Spinning Disk confocal microscope using a 60× oil-immersion objective (NA 1.40). Imaging parameters (laser power, exposure time) were held constant across all conditions within an experiment. YAP/TAZ nuclear localization was quantified as the ratio of nuclear to cytoplasmic mean fluorescence intensity using automated segmentation in ImageJ. At least 100 cells were analyzed per condition across a minimum of three biological replicates.

### Bioinformatics and computational analyses

For gene expression analysis, TCGA-BRCA RNA-seq (HTSeq-FPKM) and clinical data were accessed via the GDC portal (v36.0, acquired Jan 2023). KMPlot was also utilized to obtain clinical survival data (with parameters in supplemental data. CCLE RNA-seq (TPM) data were downloaded from the Broad Institute (2019 release, acquired Feb 2023). For copy number analysis, Genomic Identification of Significant Targets in Cancer (GISTIC) data were downloaded from cBioPortal (v6.4.4, acquired Mar 2026). For methylation analysis, Illumina 27k and 450k methylation arrays were downloaded from the GDC portal (v36.0, acquired Jan 2023). For gene expression analysis, data were log2-transformed for comparisons. PAM50 subtype assignments from the TCGA dataset were used for stratification. For copy number assessment, frequency of each alteration in TCGA-PanCan-BRCA and METABRIC datasets, stratified by PAM50 subtype, were performed. For methylation analysis, beta values were used to assess frequency of epigenetic alteration. To assess correlation of alterations and gene expression, simple linear regression analysis of both copy number or methylation status vs gene expression was performed. For survival analysis, TCGA-BRCA patients were stratified into high and low *SMPD3* expression groups based on upper and lower quartiles within each PAM50 subtype. Relapse-free survival (59) was plotted using KMPlot. The full command parameters are provided in the Supplementary Methods. For Gene Set Enrichment Analysis (GSEA v4.4.2), TCGA-BRCA patients were stratified into high and low *SMPD3* expression groups based on upper and lower deciles. Analysis was performed on a gene list ranked by the signal-to-noise ratio between *SMPD3*-high and *SMPD3*-low tumors. The MSigDB C6 oncogenic signatures collection (v7.5.1) was queried using 1,000 phenotype permutations. Significance was defined as a false discovery rate (FDR) q-value < 0.05. Full command parameters are provided in Supplementary Methods.

### Statistical analysis

Data were analyzed using GraphPad Prism (v10.3.1) and are presented as mean ± SEM from at least three independent biological replicates, unless otherwise stated. Prior to statistical testing, data distribution was assessed for normality (Shapiro-Wilk test) and homogeneity of variances (Levene’s test). Two-group comparisons were made using unpaired, two-tailed t-tests. Comparisons of more than two groups were performed using one-way or two-way ANOVA followed by Šidák’s post-hoc test for multiple comparisons. Where assumptions of normality or equal variance were not met, appropriate non-parametric tests (e.g., Mann-Whitney U test, Kruskal-Wallis test) were used. A p-value < 0.05 was considered statistically significant. Where appropriate, effect sizes (Cohen’s d or partial eta-squared) are reported. Raw data and analysis scripts will be deposited in GEO and a public GitHub repository upon publication.

### Data Availability

The TCGA-BRCA RNA-seq and clinical data analyzed in this study are available via the GDC portal (v36.0). CCLE RNA-seq data are available from the Broad Institute (2019 release). Microarray data have been deposited in the Gene Expression Omnibus under accession number [GEO: pending]. Source data for all figures are provided with this paper.

## RESULTS

### nSMase2 is suppressed and inversely associated with prognosis of basal-like and HER-2 positive BC

The functional inactivation of tumor suppressor genes is a defining hallmark of carcinogenesis (60, 61) and is typically driven through copy number loss (61), genetic mutation (61), and/or epigenetic silencing (61). The *SMPD3* gene is localized to chromosomal region 16q22.1 (43) – a common site of loss of heterozygosity in BC (47) – and we and others have shown that *SMPD3* is epigenetically suppressed in multiple cancers including BC (31, 50, 51). While this is suggestive that nSMase2 may function as a tumor suppressor within the SL network **(Fig. 1A)**, a comprehensive assessment of these alterations in BC is lacking, and the clinical consequences are unclear. To approach this, a combination of genomics, cell line studies, and clinical data was deployed to assess canonical features of tumor suppressors, including loss of heterozygosity, loss of function, and associations with patient prognosis **(Fig. 1B)**. To define the extent of genetic alterations of *SMPD3* in BC, the Genomic Identification of Significant Targets in Cancer (GISTIC) module was utilized to assess *SMPD3* copy number in BC patient cohorts from The Cancer Genome Atlas (TCGA) and Molecular Taxonomy of Breast Cancer International Consortium (METABRIC). Analysis indicated that the hemizygous deletion of *SMPD3* was observed in 61% of TCGA and 58% of METABRIC BC patients (**Fig. 1C, S1A**), a frequency comparable to established BC tumor suppressors including *PTEN* (62), *BRCA1/2* (63), and *RB1* (64). This was observed across all PAM50 molecular subtypes. To further investigate the genetic and epigenetic routes of *SMPD3* inactivation, mutational frequency and DNA methylation status (using Illumina 27k and 450k methylation arrays) were analyzed across the same patient cohorts. Notably, *SMPD3* mutations were rare (3 of 3593 BC patients) **(Fig. S1B)**. However, consistent with prior cell-based studies (65) extensive hypermethylation was observed within the *SMPD3* promoter and putative downstream enhancer regions across all BC subtypes relative to benign mammary tissue, with the highest levels detected in the HER-2 positive and basal-like tumors **(Fig. 1D)**. This suggests that epigenetic silencing is a major mechanism of *SMPD3* inactivation. Indeed, analysis of RNA-seq data from TCGA-BRCA patients revealed that *SMPD3* mRNA levels were significantly reduced in BC tumors compared to normal mammary tissue across all PAM50 subtypes (**Fig. 1E**), again consistent with those of canonical tumor suppressors in BC such as *PTEN* (66) and *RB1* (64) with *SMPD3* transcript levels exhibiting significant correlations with both copy number alterations **(Fig. S1C)** and DNA methylation status **(Fig. S1D)**. This supports both these mechanisms as contributors to gene expression loss in BC and confirms functional consequences. To assess whether *SMPD3* downregulation is preserved in cell-line model systems, *SMPD3* expression was evaluated across all available BC cell lines in the Cancer Cell Line Encyclopedia (CCLE). Consistent with patient data, all BC cell lines exhibited uniformly lower *SMPD3* expression compared to normal human mammary epithelial cells (HMECs), with basal-like and HER2-positive lines showing the greatest reductions (**Fig. 1F**). Independent validation by RT-qPCR in MCF7 (luminal), SKBR3 (HER2-positive), and MDA-MB-231 (basal-like) cells confirmed reduced *SMPD3* expression relative to the non-transformed mammary epithelial line MCF-10A (**Fig. 1G**), establishing these models as appropriate systems for interrogating the functional consequences of *SMPD3* loss. Of note, the most pronounced downregulation of *SMPD3* in cell lines and tumors was observed in basal-like and HER2-positive tumors (**Fig. 1G**), subtypes associated with the poorest clinical outcomes (67). To determine whether reduced transcript levels translate into decreased protein expression in human cancer, tumor tissue microarrays were stained for nSMase2 using a validated in-house antibody (**Fig. S1E**). Here, normal mammary ducts displayed strong, localized nSMase2 staining whereas breast tumors exhibited weaker and more diffuse staining patterns, consistent with reduced nSMase2 protein abundance (**Fig. 1H, S1F**). Finally, the clinical significance of *SMPD3* downregulation was evaluated using publicly available patient survival data (68) with patients stratified by *SMPD3* expression quartiles within each PAM50 subtype. Across all subtypes, low *SMPD3* expression was associated with significantly worse relapse-free survival with the greatest differences observed in HER2-positive and basal-like diseases (**Fig. 1I, S1G**), the same subtypes exhibiting the most profound loss of *SMPD3* expression. Collectively, these analyses demonstrate consistent and multi-faceted suppression of *SMPD3* at the genetic level in BC and link *SMPD3* loss to adverse clinical outcomes. Based on these characteristics, this supports a putative tumor suppressive role for nSMase2 in BC.

**Fig. 1.**
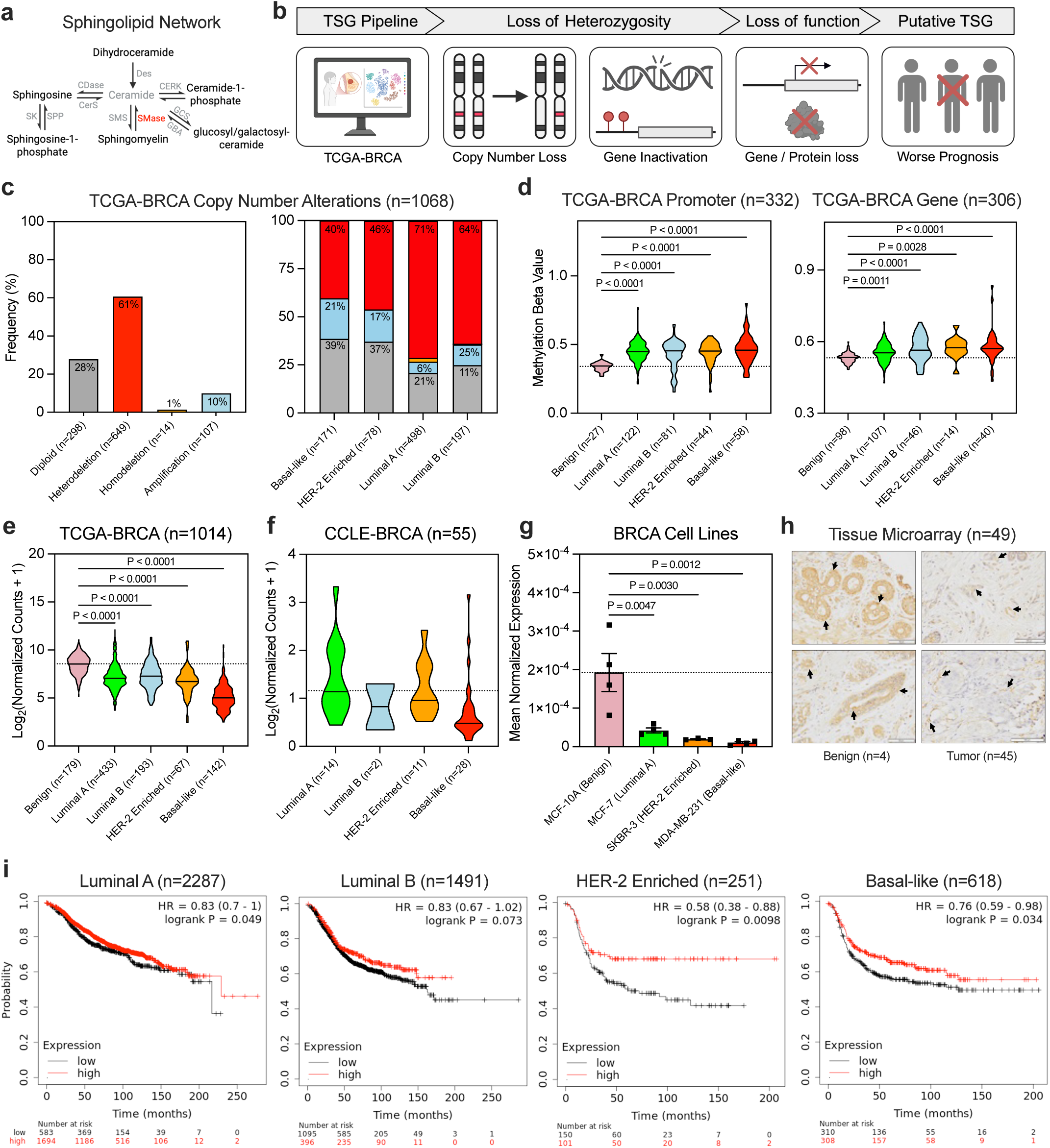
nSMase2 is suppressed and inversely associated with prognosis of basal-like and HER-2 positive BC. **a,** Condensed SL metabolic pathway; modified from Many Ceramides (19). **b,** TSG identification pipeline; TCGA, METABRIC, CCLE, tumor cohorts, and KMPLOT data sets were used to define copy number variations, gene methylation, gene expression, protein levels, and prognosis. **c,** GISTIC copy number alterations of TCGA breast tumors, stratified by PAM50 molecular subtype, compared to normal mammary epithelial tissue. **d,** Methylation beta scores of *SMPD3* gene promoter (left) and putative downstream enhancers (right) stratified by PAM50 molecular subtype. **e,** nSMase2 expression of (TCGA) breast tumors, stratified by PAM50 molecular subtype, compared to normal mammary epithelial tissue. **f,** nSMase2 expression of (CCLE) breast cancer cells, stratified by PAM50 molecular subtype, compared to normal mammary epithelial (HME1) cell line. **g,** nSMase2 expression of breast cancer cells compared to normal mammary epithelial (MCF-10A) cell line via RT-qPCR. **h,** Representative IHC staining of nSMase2 in human breast tumor and benign mammary tissue. Arrows highlighting differential expression in breast ducts. **i,** Kaplan-Meier relapse-free survival plots of breast cancer patients stratified by low vs high (quartile) nSMase2 expression of primary tumors. Gene expression in (**g)** is representative of mean ± SEM of 4 independent biological replicates unless otherwise specified. *P* values obtained via one-way ANOVA + Tukey multiple comparisons test.

### Restoration of nSMase2 activity suppresses basal-like and HER-2 positive BC tumorigenesis

To conclusively establish nSMase2 as a tumor suppressor, it became critical to assess the functional consequences of restoring nSMase2 expression. For this, lentiviral vectors were used to generate BC cell lines stably expressing V5-tagged wild-type nSMase2 (N2), a construct-matched empty vector (EV), or a catalytically inactive mutant (H639A; HA, as previously described) (69). As model cell lines, MDA-MB-231 (Basal-like) and JIMT-1 (HER2-positive) cells were selected based on low endogenous nSMase2 expression (**Fig. 1G**) and robust *in vivo* tumor-forming capacity (70, 71). Immunoblot analysis using anti-V5 and anti-nSMase2 antibodies confirmed stable overexpression of each construct (**Fig. S2A**), while immunofluorescence microscopy further confirmed plasma membrane localization of nSMase2 (**Fig. S2B**) as reported previously (69). *In vitro* nSMase activity assays revealed marked increase of enzymatic activity in N2-expressing cells relative to HA controls (**Fig. 2A**), confirming a functional restoration of nSMase2 activity. Lipidomics analysis further demonstrated that N2 but not HA cells had elevated total Cer levels (**Fig. 2B, S2C**), although notably the levels of downstream metabolites sphingosine (Sph) and sphingosine-1-phosphate (S1P) were unaltered (**Fig. S2D-E**). As EV and HA-expressing cells generated comparable Cer levels (**Fig. S2F**) and exhibited similar tumor growth *in vivo* (**Fig. S2G**), all subsequent experiments were performed with N2 and HA cells exclusively. To assess the *in vivo* effects of nSMase2 expression, cell lines were orthotopically implanted in NSG mice and monitored over time. Results indicated that restoration of nSMase2 expression did not impair tumor initiation for MDA-MB-231 and JIMT-1 cells, as both N2 and HA cell lines formed palpable and comparably sized tumors by day 7 post-implantation **(Fig. 2C)**. Similarly, early tumor growth (days 7-17) was not significantly different between groups **(Fig. 2C)**. However, beginning at day 17, tumor growth trajectories began to diverge with N2-expressing tumors exhibiting significantly reduced growth compared to HA controls **(Fig. 2C)**. Analysis of tumor size at experimental endpoint revealed a 40% (MDA-MB-231) and 46% (JIMT-1) reduction in mean tumor volume **(Fig. 2D)**. This was also reflected in the 42% (MDA-MB-231) and 26% (JIMT-1) reduction in mean tumor mass **(Fig. S2H)**. Collectively, this provides functional *in vivo* evidence that restoration of nSMase2 Cer-generating activity (**Fig. 2E, S2I-J)** exerts a tumor-suppressive effect *in vivo*.

**Fig. 2.**
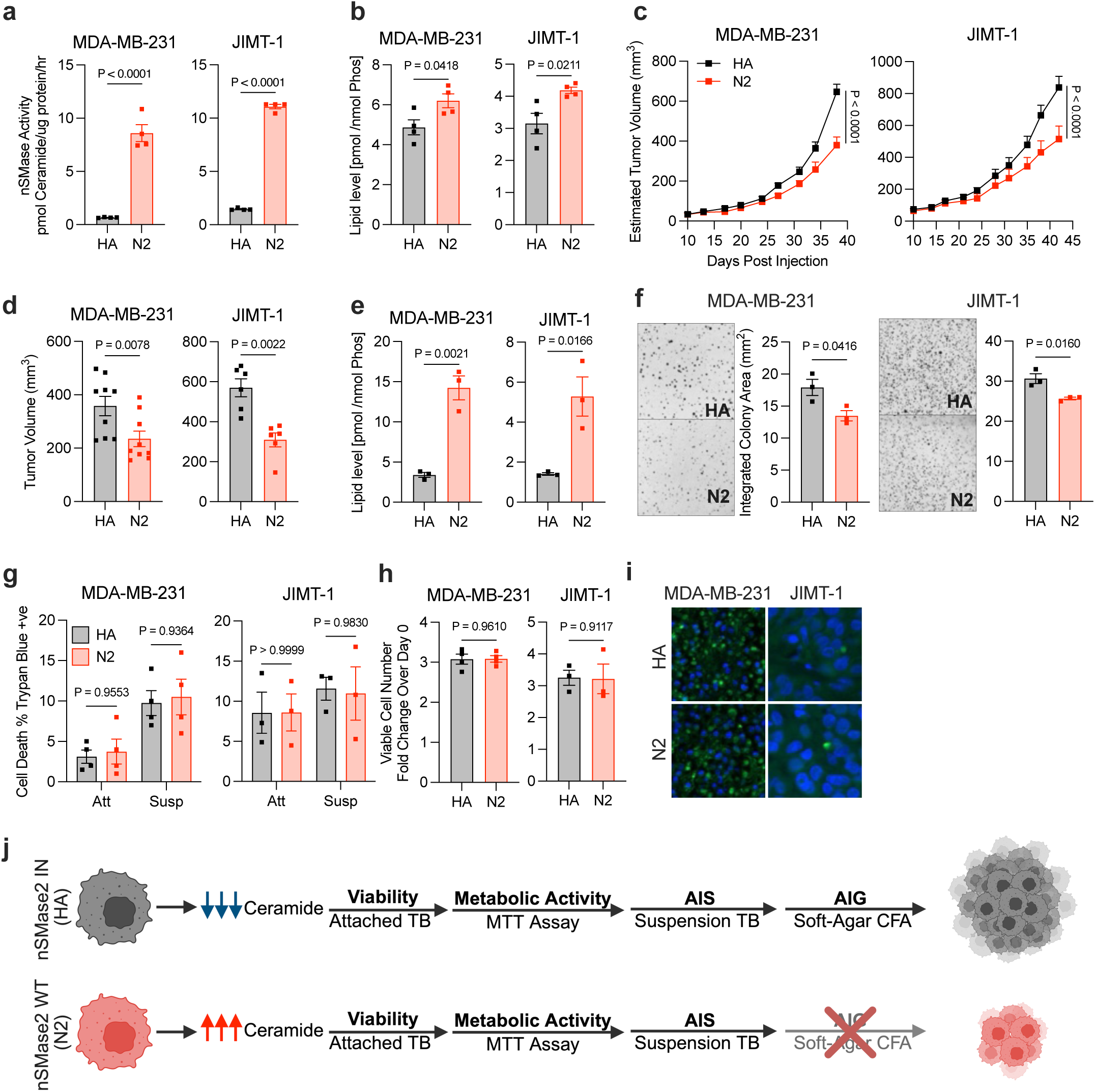
Restoration of nSMase2 activity suppresses basal-like and HER-2 positive BC tumorigenesis. **a,** MDA-MB-231 and JIMT-1 cells stably overexpressing either WT (N2) or inactive (HA) nSMase2 validated via nSMase2 radioactivity assay. **b,** Total ceramide levels of MDA-MB-231 or JIMT-1 cells via LC/MS/MS. **c,d,** MDA-MB-231 (n = 9) or JIMT-1 (n = 6) cells orthotopically implanted into the mammary fat pad of NSG mice. **c,** Tumor volume (in 2 dimensions) was measured with calipers over duration of the experiment. **d,** Tumor volume (in 3 dimensions) was measured with calipers at endpoint. **e,** Total ceramide levels of MDA-MB-231 or JIMT-1 tumors via LC/MS/MS. **f,** Soft-agar colony formation assay for 21 (JIMT-1) or 28 days (MDA-MB-231), imaged, and quantified via cell cytometer, displayed as integrated colony area. **g,** Trypan blue assay in attached and suspension conditions at 48 hrs following cell seeding. **h,** MTT viability assay of experimental samples at 72 hrs following seeding normalized to initial absorbance reading at 24 hrs. **i,** TUNEL assay of MDA-MB-231 or JIMT-1 tumors at experimental endpoint. **j,** Experimental flowchart of nSMase2 biology. **a-h,** Images are representative of mean. Data are representative of mean ± SEM of 3 or 4 independent biological replicates unless otherwise specified. *P* values obtained via student’s *t*-test or two-way ANOVA + Tukey multiple comparisons test.

Tumor suppressors typically exert their functions through alterations in cellular metabolism (72), increasing cell death (73), or inhibition of pro-growth signaling pathways (73) - with prior studies implicating nSMase2 in such biologies under various conditions (30–32, 35, 48). To begin defining the biological mechanisms underlying nSMase2-mediated tumor suppression, cells were first subjected to soft-agar colony formation assays, an established *in vitro* surrogate for cancer cell transformation and tumorigenicity. Here, nSMase2 activity significantly reduced colony formation in both MDA-MB-231 and JIMT-1 cells (**Fig. 2F, S2K**), with assay endpoints at 28 and 21 days respectively. Colony formation reflects multiple biological processes including general cell viability, anchorage-independent survival (AIS), and anchorage-independent growth (AIG), thus, additional assays were performed to isolate the precise N2-regulated mechanism. Analysis of cell viability in standard adherent conditions by MTT and trypan blue exclusion showed no significant difference between N2 and HA cells (**Fig. 2G–H, S2L-S2M**), suggesting that nSMase2 does not exert a significant effect on cell death. Consistent with this, TUNEL staining of tumor sections revealed no significant differences in TUNEL positivity between N2 and HA tumors (**Fig. 2I**), and assessment of AIS by forced suspension cultures in low-attachment trays similarly revealed no significant differences between N2 and HA cells (**Fig. 2G, S2M**). Taken together, this supports a model in which the nSMase2–Cer axis selectively suppresses AIG (**Fig. 2J**).

Given that AIG is a key determinant of tumor aggressiveness (74), effects of N2 expression on spontaneous metastasis were evaluated. Here, histological analysis of lung sections revealed a marked reduction in metastatic burden in mice bearing N2-expressing tumors compared to HA controls (**Fig. S2N**). However, this was not due to impaired motility as migration capacity in transwell assays was comparable between N2- and HA-expressing cells (**Fig. S2O**), suggesting that reduced metastatic burden is likely a secondary consequence of diminished primary tumor growth. Collectively, these data demonstrate that restoration of nSMase2 activity suppresses breast tumor growth and metastasis primarily through effects on AIG, a defining hallmark of malignant progression (**Fig. 2J**).

### TAZ identified as a major downstream effector of nSMase2

Despite its importance as a central hallmark of cancer transformation, the molecular mechanisms governing the acquisition of AIG remain poorly defined. To begin to elucidate the mechanisms by which nSMase2 suppresses AIG and tumor growth, microarray analysis was performed on RNA extracted from MDA-MB-231 N2 and HA tumors. Gene-expression analysis revealed surprisingly modest transcriptional changes in N2 tumors with 11 genes significantly upregulated and 29 genes significantly downregulated compared to HA controls (FDR < 0.05; > 2-fold change) (**Fig. S3A**). To identify the biological implications of these alterations, pathway enrichment analysis was performed on the differentially expressed gene set. This identified YAP1–extracellular matrix (ECM) signaling as the most significantly altered pathway in nSMase2-expressing tumors (**Fig. 3A**). RT-qPCR analysis of an independent cohort of xenografted tumors confirmed microarray results showing that N2 tumors exhibited a pronounced reduction in established downstream YAP/TAZ target genes (75, 76) including *EDNRA*, *THBS2*, and *GPC6* (**Fig. 3B**) and a modest but significant suppression of YAP and TAZ expression (**Fig. 3B**), without changes in gene expression of the binding partner TEAD2 or other YAP/TAZ related genes (**Fig. S3C)**. To complement the tumor-based analysis, an orthogonal clinical dataset approach was employed to identify pathways linked with nSMase2 expression in patient tumors. Gene-set enrichment analysis (GSEA) was performed using TCGA breast cancer datasets stratified by nSMase2 expression deciles. While multiple biological pathways were identified, nSMase2-low tumors (n = 147) were strikingly enriched for a YAP1-Up gene signature, whereas nSMase2-high tumors (n = 134) were enriched for a YAP1-Down signature (**Fig. 3C**). Consistently, Integrated Pathway Analysis demonstrated significantly reduced YAP and TAZ pathway activity in nSMase2-high tumors (**Fig. S3C**). Together these analyses establish an inverse relationship between nSMase2 expression and YAP/TAZ signaling in cell-line derived and patient tumors. The results also suggest that the YAP/TAZ pathway is a downstream target of the tumor suppressive functions of nSMase2.

**Fig. 3.**
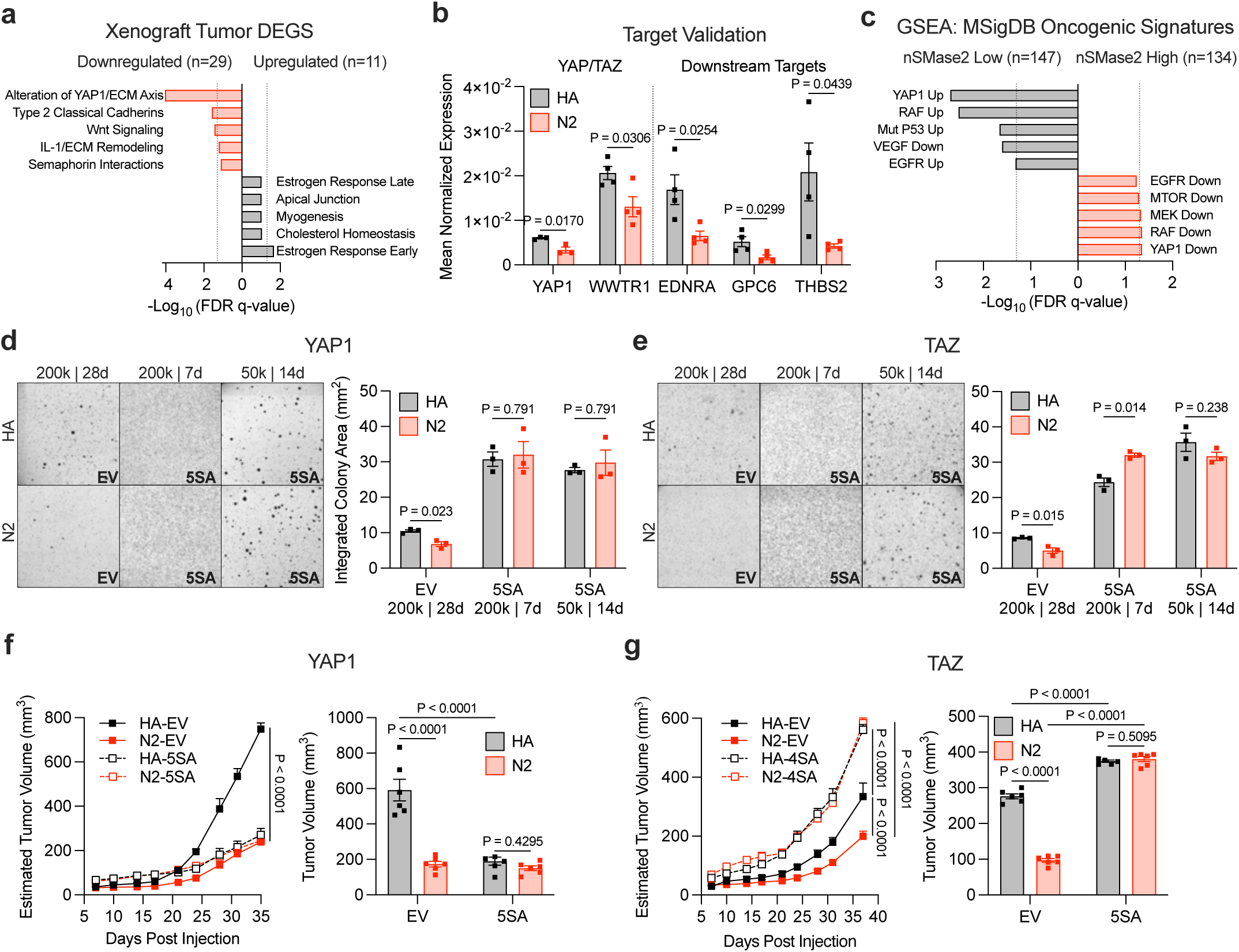
TAZ identified as a major downstream effector of nSMase2. **a,** Pathway enrichment of genechip microarray of MDA-MB-231 xenografted tumors (n = 4). **b,** RT-qPCR validation of YAP/TAZ and downstream targets identified in microarray in MDA-MB-231 tumors (n = 4). **c,** Gene set enrichment analysis of cohort of breast cancer patients (TCGA) stratified by (quartile) nSMase2 expression. **d,e,** Soft-agar colony formation assay of MDA-MB-231 cells overexpressing **d,** YAP1 or **e,** TAZ, imaged, and quantified via cell cytometer, displayed as integrated colony area. **f,g** MDA-MB-231 cells (n = 6) overexpressing **f,** YAP1 or **g,** TAZ orthotopically implanted into the mammary fat pad of NSG mice. Tumor volume (in 2 dimensions) was measured with calipers over duration of the experiment. Tumor volume (in 3 dimensions) was measured with calipers at endpoint. **b,d-g** Data are representative of mean ± SEM of 3 or 4 independent biological replicates unless otherwise specified. *P* values obtained via student’s *t*-test or two-way ANOVA + Tukey multiple comparisons test.

Having identified YAP/TAZ as potential target of nSMase2, it was crucial to establish if suppression of YAP/TAZ signaling is functionally required for the anti-tumor activity of nSMase2. For this, lentiviral transduction was used to introduce previously reported YAP-5SA, TAZ-4SA mutants, or an empty vector (EV) control into HA- and N2-expressing cells (**Fig. S3D**). These widely used mutants (77, 78) are resistant to inhibitory phosphorylation signals, resulting in their constitutive nuclear localization and impaired degradation; thus, enhancing their activity. Immunoblot analysis confirmed robust expression of each construct (**Fig. S3E**), and immunofluorescence microscopy demonstrated strong *in vitro* (**Fig. S3F**) *and in vivo* (**Fig. S3G-H**) nuclear localization irrespective of nSMase2 status consistent with enforced pathway activation. *In vitro* soft agar assays demonstrated that expression of either YAP-5SA or TAZ-4SA alone dramatically enhanced colony formation in HA cells (**Fig. 3D-E**), consistent with prior reports identifying both YAP and TAZ as key drivers of AIG. Importantly, reactivation of either paralog was sufficient to overcome nSMase2-mediated suppression of colony formation (**Fig. 3D-E**), demonstrating a key role for inhibition of the YAP/TAZ pathway in mediating the AIG-suppressive effects of nSMase2.

To extend these findings *in vivo*, HA- and N2-expressing cells transduced with YAP-5SA, TAZ-4SA or their respective EV controls were orthotopically implanted into NSG mice. Importantly, for both EV control cell lines, earlier results **(Fig. 2C-D)** were recapitulated with nSMase2 activity leading to suppression of primary tumor growth compared to HA controls (**Fig. 3F**). Strikingly, expression of YAP-5SA led to earlier tumor initiation (palpable tumors at 4–5 days versus 7 days in EV controls) and increased initial tumor size in both HA and N2 backgrounds (**Fig. S3I**). However, by day 17, tumor-growth trajectories diverged with YAP-5SA expression in both N2 and HA tumors leading to a significant growth suppression to levels comparable to N2-EV tumors (**Fig. 3F**), demonstrating a biphasic action of YAP on tumor growth. Expression of TAZ-4SA also led to earlier tumor initiation (palpable tumors at 4–5 days versus 7 days in EV controls) and increased initial tumor size in both HA and N2 backgrounds (**Fig. 3G, S3J**). However, unlike YAP, these tumor-growth trajectories continued, and TAZ-4SA resulted in robust tumor growth in both HA and N2 cells, with no significant difference between the two groups (**Fig. 3G, S3J**). These findings demonstrate that reactivation of TAZ, but not YAP, is sufficient to bypass nSMase2-mediated suppression of tumorigenesis. Taken together, these findings demonstrate that the tumor-suppressive effects of nSMase2 are mediated through selective inhibition of a TAZ-dependent, but not YAP-dependent program. Moreover, these results suggest that TAZ, rather than YAP, functions as the dominant oncogenic Hippo effector driving late-stage breast cancer progression.

### nSMase2–Cer axis suppresses TAZ nuclear localization

TAZ is a transcriptional co-activator that exerts its activity upon binding to transcription factor partners within the nucleus. Given that nSMase2 expression led to suppression of YAP/TAZ-regulated genes, the effects of nSMase2 on nuclear localization of TAZ were evaluated. To assess this *in vivo*, immunofluorescence analysis of N2 and HA tumors was performed, and as can be seen, in tumors at day 17 post-implantation – which is typically prior to the time point at which N2 and HA tumor growth diverges (**Fig. 2C**), TAZ was predominantly nuclear in both HA- and N2-expressing tumors, whereas YAP remained largely cytoplasmic, suggesting it is transcriptionally inactive (**Fig. S4A**). Analysis of tumors at experimental endpoints – where nSMase2 suppression of growth is maximal (**Fig. 2C**) – showed a marked reduction of nuclear TAZ in N2 tumors but not in HA tumors (**Fig. 4A**). In contrast, YAP localization was unchanged between N2 and HA tumors, showing a predominantly cytosolic localization (**Fig. 4B**). Together, these findings suggest that YAP is largely inactive *in vivo* while TAZ is nuclear localized and transcriptionally active at both early and late stages of tumor progression. Moreover, these results indicate that an active nSMase2–ceramide (Cer) axis selectively suppresses nuclear localization of TAZ to exert its tumor suppressive effects.

**Fig. 4.**
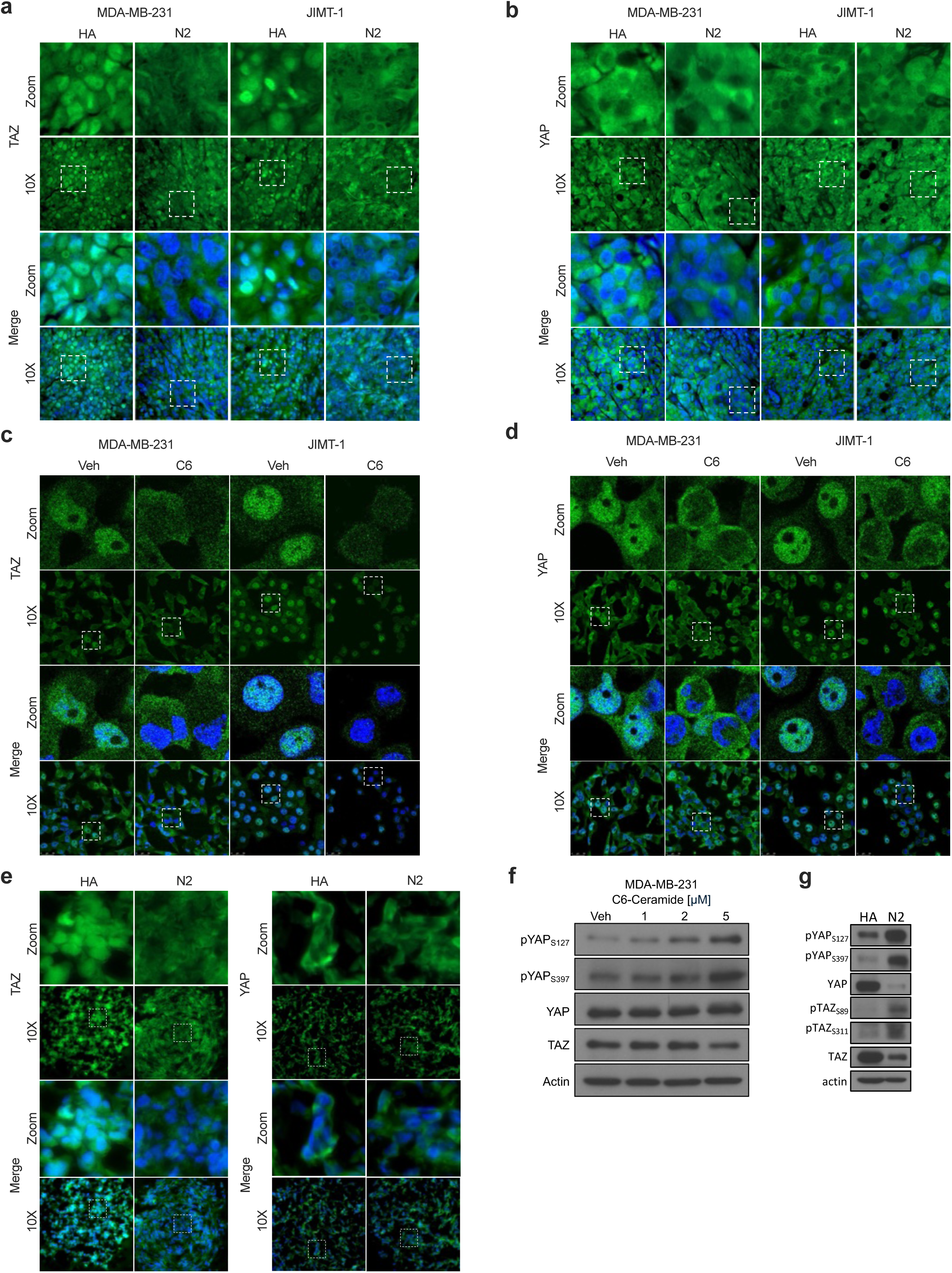
nSMase2–Cer axis suppresses TAZ nuclear localization. **a-e,** Assessment of YAP/TAZ localization via immunofluorescent microscopy with anti-YAP and anti-TAZ antibodies with DAPI used as nuclear co-stain. Quantification of nuclear localized YAP/TAZ was performed manually via blinded observer. **a-b,** MDA-MB-231 and JIMT-1 tumors (n = 3). **c-d,** MDA-MB-231 and JIMT-1 cells (n = 3) in monolayer treated with either C6-Cer or Veh control. **e,** 3D spheroids of MDA-MB-231 cells. **f,** Assessment of the indicated protein and phosphoprotein levels in MDA-MB-231 and JIMT-1 cells (n = 3) in monolayer treated with either C6-Cer or Veh control via immunoblot. **g,** Assessment of the indicated protein and phosphoprotein levels in MDA-MB-231 spheroids via immunoblot. **a-e,** Data are representative of mean ± SEM of 3 or 4 independent biological replicates unless otherwise specified. *P* values obtained via student’s *t*-test.

The suppression of YAP and TAZ nuclear localization can occur through either cell-intrinsic mechanisms or tissue specific mechano-transduction machinery. In order to further investigate the context by which nSMase2 regulates the YAP/TAZ pathway, it was first necessary to determine if these effects are specific to the complex *in vivo* tumor environment. Immunofluorescence analysis in standard monolayer conditions showed that both YAP and TAZ were localized to the nucleus in the majority of cells regardless of nSMase2 activity status (**Fig. S4B-C**). Moreover, nSMase2 expression did not alter transcript levels of downstream TAZ target genes in neither adherent conditions nor in short-term suspension cultures (**Fig. S4D**). Prior studies in other systems have reported that exogenous Cer suppresses YAP/TAZ signaling and localization in monolayer conditions. Here, treatment of MDA-MB-231 and JIMT-1 cells with exogenous C6-ceramide caused a similar suppression of both YAP and TAZ nuclear localization in monolayer culture (**Fig. 4C-D**). This confirms that a generalized increase in Cer signaling is sufficient to inhibit the Hippo pathway in monolayer culture but that the nSMase2-mediated suppression of TAZ occurs specifically within three-dimensional contexts associated with AIG and *in vivo* tumor biology. This may also account for why nSMase2 expression had no effect on monolayer growth (**Fig. 2G-H**). To more accurately model the three-dimensional tumor environment in which nSMase2-derived Cer exerts its effects, YAP and TAZ localization was examined in 3D spheroid culture. Here, YAP was found to be cytoplasmic in both HA and N2 cells while TAZ was predominantly nuclear in HA spheroids and was retained in the cytoplasm in N2 spheroids (**Fig. 4E**). Together, these findings demonstrate that nSMase2 suppresses tumorigenesis through selective, context-dependent inhibition of TAZ nuclear localization. They also suggest that exogenous Cer treatment in monolayer culture and 3D spheroid systems are tractable models for dissecting the molecular mechanisms downstream of nSMase2-Cer signaling.

The suppression of cell-intrinsic YAP and TAZ nuclear localization can occur through a variety of pathways including but not limited to activation of the Hippo kinase core (79), subsequent degradation via ubiquitin-mediated E3 ligases (80), and upstream signal transduction of focal adhesion kinases (81). To define the molecular mechanism by which the nSMase2–Cer axis suppresses TAZ nuclear localization, acutely probe the signaling pathway, and identify the maximal effects of Cer signaling, we initially utilized exogenous C6-Cer. C6-Cer treatment (5 µM) induced the phosphorylation of YAP at serine residues S127 and S397 within 4 h with a maximal phosphorylation detected 8 h post-treatment while by 12 h of treatment, both YAP and TAZ protein levels were decreasing (**Fig. S4E**). This suggests that Cer induces phosphorylation of YAP at the LATS kinase consensus site S127 followed by phosphorylation at S397, which primes YAP/TAZ for proteasomal degradation. Of note, in these studies, direct detection of TAZ phosphorylation was not feasible using the available antibodies. To assess the Cer–Hippo signaling axis at the lowest effective signaling threshold, a C6-Cer dose response was performed at 8h with results showing dose-dependent increases in YAP phosphorylation at both S127 and S397 (**Fig. 4F, S4F**). Notably, C6-Cer did not alter MST1/2 phosphorylation levels but modestly increased the p-LATS/LATS ratio **(Fig. S4G)** suggesting that LATS kinase is the principal Cer-regulated node within the Hippo pathway. Furthermore, neither the inactive L-erythro-C6-Cer isomer nor other sphingolipid metabolites (dhCer, dhSph, Sph) were effective at promoting YAP phosphorylation at either site (**Fig. S4H**), suggesting that this is a Cer-specific regulation. Importantly, these results were recapitulated in 3D spheroid cultures of N2- and HA-expressing cells (**Fig. 4G**), reinforcing that the action of nSMase2 is through Cer. Together, these findings demonstrate that the nSMase2-Cer axis suppresses TAZ nuclear localization with LATS kinase as the most proximal signaling node.

### LATS kinase is required for tumor suppressive functions of nSMase2

Having linked LATS kinase to nSMase2, it became critical to directly test if LATS kinase is necessary for Cer-mediated YAP/TAZ phosphorylation. Here, cells were pre-treated with the LATS kinase inhibitor TRULI (5 µM, 12 h) at the minimal effective dose required to suppress YAP phosphorylation under both adherent and suspension conditions (**Fig. S5A**) and then treated with C6-Cer (5 µM, 8 h). Pre-treatment with LATS inhibitor TRULI markedly attenuated both the phosphorylation of YAP (S397) and TAZ (S89 and S311) (**Fig. 5A)** and Cer-dependent suppression of nuclear localization (**Fig. 5B-C)**, suggesting that the Cer-mediated effects on YAP/TAZ are LATS kinase dependent. To determine if N2-derived Cer signaling also follows this paradigm, 3D spheroid cultures of N2- and HA-expressing cells were treated with TRULI (5 µM, 48 h) or a vehicle control. Here, nSMase2 activity suppressed TAZ but not YAP nuclear localization in spheroids, and this suppression did not occur in spheroids treated with TRULI (**Fig. 5D-E, S5B)**, suggesting that the nSMase2-derived Cer effect on YAP/TAZ is also LATS dependent. Finally, the effect of pharmacologic LATS inhibition on soft-agar colony formation was performed, and the results showed that LATS inhibition completely restored AIG in N2-expressing cells (**Fig. 5F**). Together, these findings demonstrate that nSMase2–Cer regulates the YAP/TAZ pathway through a LATS-dependent mechanism, and this leads to suppression of AIG. Collectively, these data establish the nSMase2–ceramide–LATS signaling axis as a central mechanistic pathway through which nSMase2 exerts its tumor-suppressive function (**Fig. 5G**).

**Fig. 5.**
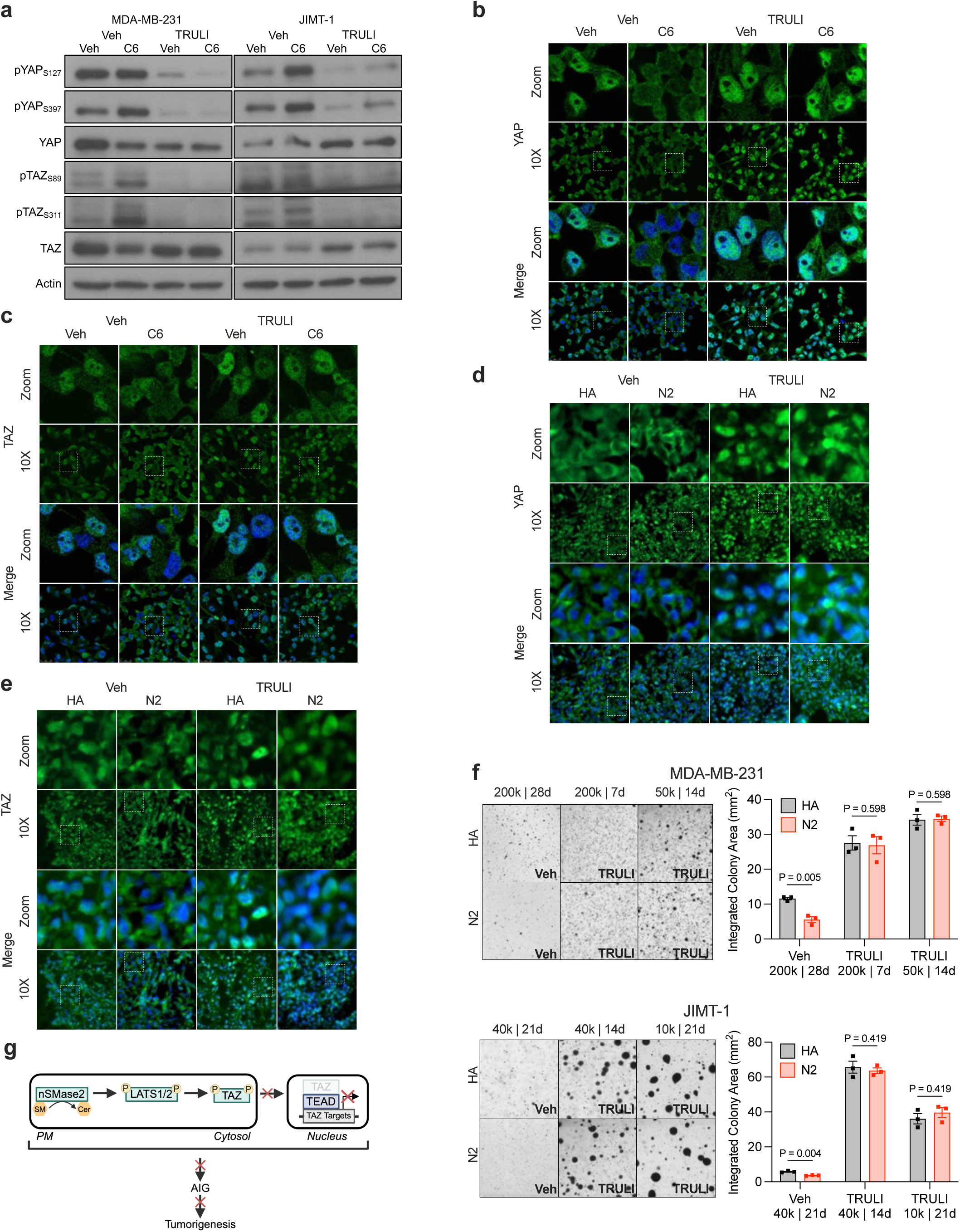
Tumor suppressive effects of nSMase2 require LATS kinase activity. **a,** Assessment of the indicated protein and phosphoprotein levels in MDA-MB-231 and JIMT-1 cells (n = 3) in monolayer treated first with TRULI (5 μM, 4 h) then with either C6-Cer (5 μM, 8 h) or Veh control via immunoblot. **b-c,** Assessment of localization of YAP and TAZ in MDA-MB-231 cells (n = 3) in monolayer treated first with TRULI (5 μM, 4 h) then with either C6-Cer (5 μM, 12 h) or Veh control via immunoblot. **d-e,** Assessment of localization of YAP and TAZ in MDA-MB-231 spheroids cultured for 12 days then treated with TRULI (5 μM, 48 h). **f,** Soft-agar colony formation assay of MDA-MB-231 and JIMT-1 cells with +/- TRULI, imaged, and quantified via cell cytometer. **g,** Schematic of proposed N2-Cer-TAZ model. **b-e,** Data is representative of mean ± SEM of 3 or 4 independent biological replicates unless otherwise specified. *P* values obtained via student’s *t*-test or two-way ANOVA + Tukey multiple comparisons test.

## DISCUSSION

While the therapeutic restoration of TSG signaling has been considered as a potential approach for treatment of aggressive cancers, clinical success has been limited owing to issues of specificity, context dependence, and druggability (17–18). Here, we have explored SL metabolism as an alternative framework for reactivating tumor suppressive programs based on the long-standing hypothesis of Cer as a candidate tumor suppressor lipid. We identify nSMase2 as a key metabolic node that is often lost or silenced in BC, and whose restoration restrains AIG in aggressive BC growth through inhibition of TAZ-dependent signaling. Taken together, these findings define a previously unrecognized tumor suppressor pathway, sheds new mechanistic light on how dysregulation of SLs drives tumorigenesis, and point to strategies aimed at restoring nSMase2-dependent Cer signaling as a viable therapeutic option.

Cer has long been considered the prototypical anti-tumor SL owing to its roles in cell death, senescence, differentiation, and growth arrest (24–27). Despite this, the acceptance of Cer as a tumor suppressor lipid has been hampered by deficiencies in our understanding of its mechanisms of generation and its specific functions, issues arising from the multiplicity of Cer-mediated pathways (19, 28–35). Major advances from this study emerged from a focus on one critical pathway of Cer generation, mediated by nSMase2. The results establish nSMase2 as a novel and *bona fide* tumor suppressor in BC through multiple lines of evidence. Bioinformatic analysis of patient data revealed significant copy number loss, and, while mutations in nSMase2 are rare, the SMPD3 gene is subjected to extensive epigenetic silencing in aggressive disease. Crucially, the magnitude and frequency of *SMPD3* suppression is consistent with functional loss of both alleles as described by Knudson’s two-hit hypothesis (7) while the strong association with poor clinical outcomes parallels what is observed for established BC tumor suppressors including TP53, PTEN, and RB1 (4–6). Biologically, the restoration of nSMase2 in aggressive BC cells led to robust suppression of *in vivo* tumor growth and restrained *in vitro* AIG. Furthermore, the selective constrainment of AIG by nSMase2 suggests that it does not function as a classical gatekeeper TSG that initiates oncogenesis, but instead resembles a “caretaker” TSG whose loss facilitates tumor progression by permitting escape from growth-limiting constraints as does BRCA1/2 loss (63). This is supported by prior studies in nSMase2-activity null mice showing spontaneous development of liver tumors but only in mice of 2 years of age or older (48). The reported epigenetic suppression of nSMase2 across a number of different cancers and pre-cancerous lesions (31, 50, 51, 65) further suggests that the potential of nSMase2 as a TSG could extend beyond BC. Of note, while results clearly establish that nSMase2 is effective at suppressing tumor intrinsic growth, the possibility of tumor extrinsic effects of nSMase2 should not be overlooked – particularly as nSMase2 is a well-established regulator of extracellular vesicle (EV) function. While a prior study in BC reported that nSMase2-dependent EV secretion led to increased angiogenesis and metastasis (56), another study showed that nSMase2-dependent EV secretion led to an enhanced response to anti-PD-1 immunotherapy (49). These reports suggest additional immune modulatory roles for nSMase2 in tumorigenesis, and this is the subject of ongoing studies in our laboratory.

Mechanistically, this study implicates TAZ a key mediator of the biologic functions of nSMase2 in tumor suppression. TAZ and YAP are paralogs and key downstream effectors of the Hippo signaling pathway that have been linked with numerous pro-tumor biologies including EMT, cell cycle progression, and AIG (77–81). Mechanistically, nSMase2-Cer inhibited TAZ functions in tumors *in vivo,* and regulated TAZ phosphorylation and levels in cells as well as and *in vivo*, resulting in the prevention of nuclear localization of TAZ. The results moreover implicate the LATS kinase as the proximal signaling node coupling TAZ to nSMase2-Cer. These findings are consistent with earlier studies linking Cer to Hippo signaling in hepatic stellate cells (82) and BC stem-like populations (83). However, a key feature of the current study is that the signaling and biological effects of the nSMase2–Hippo axis were only observed in the 3D and AIG context and not in monolayer culture. Exogenous Cer treatment in monolayer was able to mimic the signaling effects of nSMase2 on YAP/TAZ in spheroids. This raises the possibility that Cer functions as a brake on the YAP/TAZ pathway but the enzymatic Cer source varies according to the specific environmental contexts. It should also be noted that while we cannot formally exclude contributions from downstream metabolites of nSMase2-derived Cer in this study, we did not observe alterations in the levels of Sph or S1P in our nSMase2-expressing cells either *in vitro* or *in vivo*.

Importantly, this study uncovered a previously unappreciated paralog-specific divergence within the Hippo pathway. While YAP and TAZ are frequently treated as functionally equivalent transcriptional co-activators, the results here show that TAZ remained nuclear and active throughout progression of control tumors, whereas YAP was primarily cytoplasmic and likely inactive across tumor stages. This was further illustrated by the divergent effects of forced expression of active TAZ and YAP on primary tumor growth. While both YAP and TAZ were effective at promoting tumor initiation and growth in the early stages, active YAP ultimately led to restraints on tumor growth at late stages while sustained TAZ activation promoted late-stage tumor growth and overcame the effects of N2 suppression. Altogether, this suggests TAZ is the dominant oncogenic effector sustaining late-stage BC growth and supports a model of YAP as a tumor suppressor in BC, which has previously been appreciated in ER+ BCs (84). This could have important therapeutic implications as current strategies targeting YAP/TAZ signaling inhibit both paralogs by disrupting TEAD binding (85) – which could limit efficacy through antagonistic effects. In contrast, these results offer modulation of nSMase2-dependent Cer signaling as a means to selectively restore endogenous constraints on TAZ activity to restrain tumorigenesis.

In conclusion, these findings establish nSMase2 as a metabolic tumor suppressor that restrains BC growth through inhibition of TAZ signaling. This work clarifies longstanding ambiguities in Cer biology, defines novel effectors of nSMase2, and suggests divergent functions of TAZ and YAP in driving BC tumorigenesis. Reinstating these suppressive constraints by modulating nSMase2-derived Cer pools could provide a strategy for targeting aggressive BCs that are refractory to conventional oncogene-directed therapies. Conversely, loss of nSMase2 could open up opportunities to target synthetic-lethal vulnerabilities similarly to the use of PARP inhibitors in BRCA1/2-deficient tumors.

## RESOURCE AVAILABILITY

Requests for further information, resources, and reagents should be directed to and will be fulfilled by the corresponding author(s), upon request. Raw data reported in this paper is reported in Source Data. Computer code will be publicly available at GitHub: TBD. Any additional information required to reanalyze the data in this paper is available from the corresponding author(s), upon request.

## ACKNOWLEDGEMENTS

The authors would like to acknowledge the technical support provided by the Research Histology Core Laboratory - Department of Pathology, the Cancer Center Lipidomics Shared Resource, the Stony Brook Genomics Core, and the Division of Laboratory Animal Resources at Stony Brook University. We are also grateful to all members of the Lipid Cancer lab at Stony Brook University for helpful feedback and discussion throughout the duration of this study.

## AUTHOR CONTRIBUTIONS

Conceptualization, A.E.R., Y.A.H., and C.J.C.; methodology, A.E.R., I.D.M., F.N.V., M.D., Y.A.H., and C.J.C.; formal analysis, A.E.R., S.B.C., Y.A.H., and C.J.C.; investigation, A.E.R., V.F., B.K.G., S.B.C., S.L., M.E.A., D.M.P., J.O., I.P., N.C., F.N.V.; writing – original draft, A.E.R., C.J.C.; writing – review and editing, A.E.R., Y.A.H., and C.J.C.; supervision, Y.A.H. and C.J.C.; funding acquisition, M.D., Y.A.H., and C.J.C.

