## Supplemental Information for "Neutral Sphingomyelinase-2 Restrains TAZ to Suppress Breast Tumor Growth"

### FIGURE S1

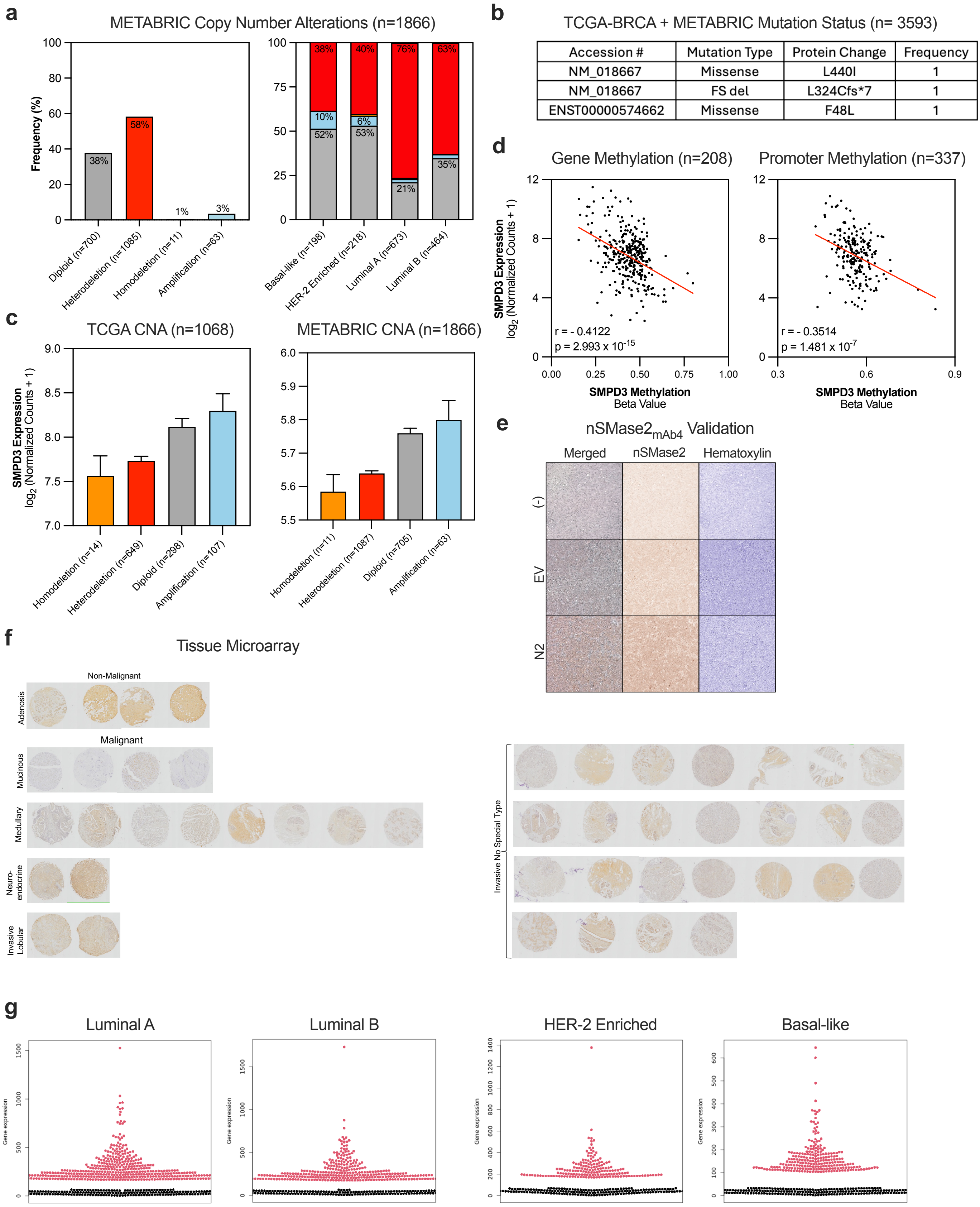


| Accession # | Mutation Type | Protein Change | Frequency |
|---|---|---|---|
| NM_018667 | Missense | L440I | 1 |
| NM_018667 | FS del | L324Cfs*7 | 1 |
| ENST00000574662 | Missense | F48L | 1 |

**Fig. S1 | nSMase2 is suppressed and inversely associated with prognosis of basal-like and HER-2 positive BC. a,** GISTIC copy number alterations (CNA) of METABRIC breast tumors, stratified by PAM50 molecular subtype, compared to normal mammary epithelial tissue. **b,** Mutation status of nSMase2 in all TCGA-BRCA and METABRIC BC patient tumor samples. **c,** nSMase2 RNA expression vs CNA in TCGA and METABRIC databases. **d,** Linear regression model via Pearson correlation of nSMase2 methylation and expression. **e,** Validation of in-house nSMase2 monoclonal antibody via IHC or immunoblot of JIMT-1 EV or N2 tumors (n = 3). IHC counterstained with hematoxylin and with deconvoluted into individual channels. **f,** Tissue microarray of human breast tumors with nSMase2 antibody. **g,** Beeswarm plots of Kaplan-Meier relapse-free survival plots. *P* values obtained via one-way ANOVA + Tukey multiple comparisons test.

### FIGURE S2

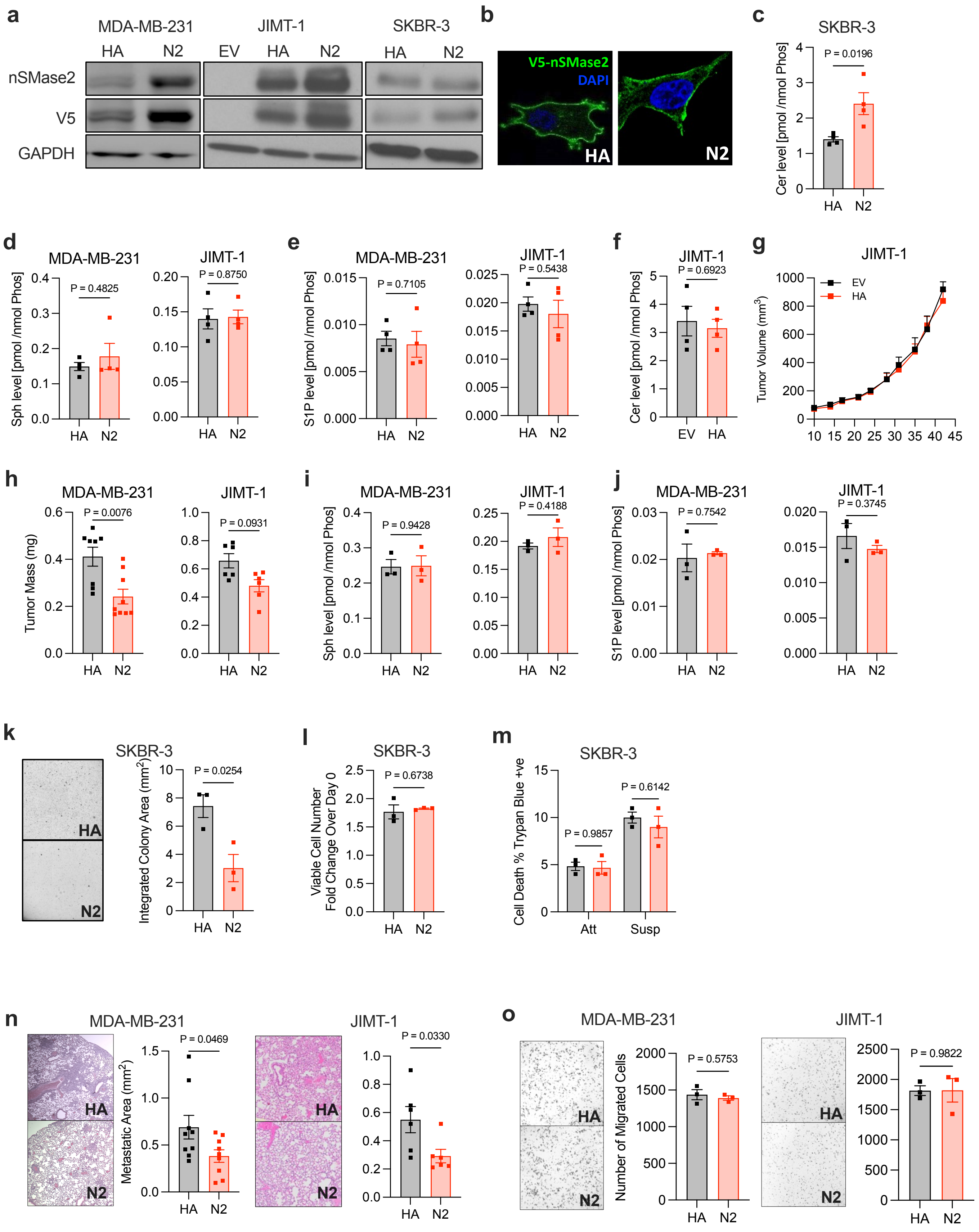

a
MDA-MB-231
JIMT-1
SKBR-3
HA
N2
EV
nSMase2
V5
GAPDH
b
V5-nSMase2
DAPI
HA
N2
c
SKBR-3
Cer level [pmol /nmol Phos]
P = 0.0196
d
MDA-MB-231
JIMT-1
Sph level [pmol /nmol Phos]
P = 0.4825
P = 0.8750
e
S1P level [pmol /nmol Phos]
P = 0.7105
P = 0.5438
f
P = 0.6923
g
Tumor Volume (mm³)
EV
HA
h
Tumor Mass (mg)
P = 0.0076
P = 0.0931
i
P = 0.9428
P = 0.4188
j
P = 0.7542
P = 0.3745
k
Integrated Colony Area (mm²)
P = 0.0254
l
Viable Cell Number
Fold Change Over Day 0
P = 0.6738
m
Cell Death % Trypan Blue +ve
P = 0.9857
P = 0.6142
Att
Susp
n
Metastatic Area (mm²)
P = 0.0469
P = 0.0330
o
Number of Migrated Cells
P = 0.5753
P = 0.9822

**Fig. S2 | Restoration of nSMase2 activity suppresses basal-like and HER-2 positive BC tumorigenesis. a,** MDA-MB-231, JIMT-1, and SKBR-3 cells stably overexpressing V5-tagged WT (N2) or inactive (HA) nSMase2 validated via immunoblot. **b,** Assessment of nSMase2 localization in MDA-MB-231 cells via immunofluorescent microscopy. **c,** Total ceramide levels of SKBR-3 cells via LC/MS/MS. **d,** Sph levels of MDA-MB-231 and JIMT-1 cells via LC/MS/MS. **e,** S1P levels of MDA-MB-231 and JIMT-1 cells via LC/MS/MS. **f,** Total ceramide levels of JIMT-1 EV or HA cells via LC/MS/MS. **g,** JIMT-1 (n = 6) cells orthotopically implanted into the mammary fat pad of NSG mice. Tumor volume (in 2 dimensions) was measured with calipers over duration of the experiment. Of note, this is the same experiment as shown in Fig. 2 but demonstrating the EV vs HA comparison. **h,** Tumor mass of MDA-MB-231 or JIMT-1 xenografts expressing HA or wild type N2. **i,** Sph levels of MDA-MB-231 and JIMT-1 tumors via LC/MS/MS. **j,** S1P levels of MDA-MB-231 and JIMT-1 tumors via LC/MS/MS. **k,** Soft-agar colony formation assay of SKBR-3 cells expressing HA or wild type N2 imaged, and quantified via cell cytometer, displayed as integrated colony area. **l,** MTT viability assay of experimental samples at 72 hrs following seeding normalized to initial absorbance reading at 24 hrs. **m,** Trypan blue assay in attached and suspension conditions at 48 hrs following cell seeding. **n,** Extracted lungs from HA and N2 injected mice were FFPE and processed for H&E; metastatic burden was assessed via ImageJ. **o,** Transwell migration assay of HA or N2, MDA-MB-231 or JIMT-1 cells for 24 hrs quantified via ImageJ. **c-o,** Data is representative of mean ± SEM of 3 or 4 independent biological replicates unless otherwise specified. *P* values obtained via student's *t*-test or two-way ANOVA + Tukey multiple comparisons test.

**FIGURE S3**

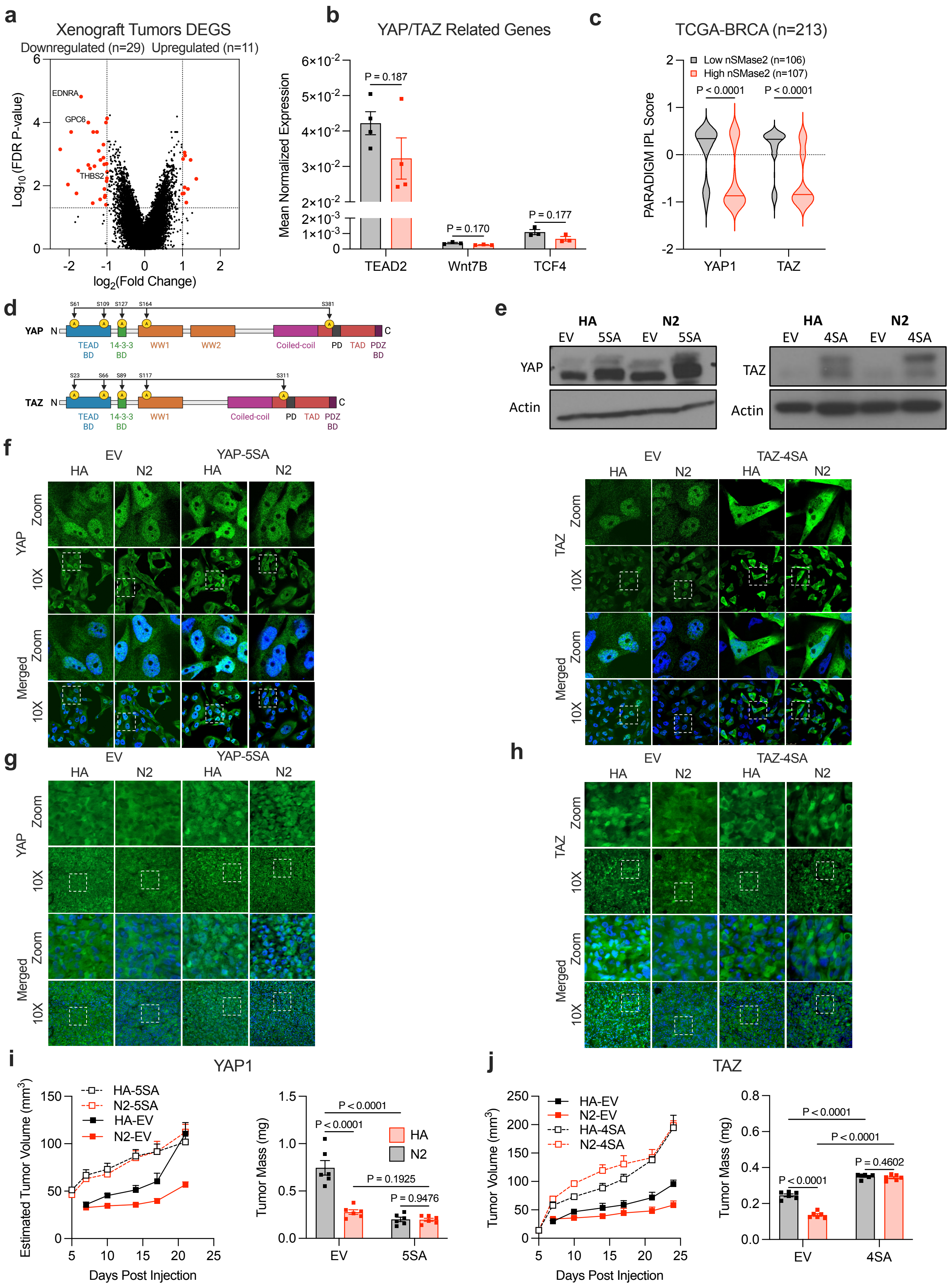

a
Xenograft Tumors DEGS
Downregulated (n=29) Upregulated (n=11)
EDNRA
GPC6
THBS2
Log10 (FDR P-value)
log2(Fold Change)
b
YAP/TAZ Related Genes
Mean Normalized Expression
P = 0.187
P = 0.170
P = 0.177
TEAD2
Wnt7B
TCF4
c
TCGA-BRCA (n=213)
Low nSMase2 (n=106)
High nSMase2 (n=107)
P < 0.0001
P < 0.0001
PARADIGM IPL Score
YAP1
TAZ
d
YAP
TAZ
TEAD BD
14-3-3
WW1
WW2
Coiled-coil
PD
TAD
PDZ BD
e
HA
N2
EV
5SA
4SA
YAP
TAZ
Actin
f
EV
YAP-5SA
TAZ-4SA
Zoom
10X
Merged
g
h
i
YAP1
HA-5SA
N2-5SA
HA-EV
N2-EV
Estimated Tumor Volume (mm3)
Days Post Injection
Tumor Mass (mg)
P < 0.0001
P = 0.1925
P = 0.9476
j
TAZ
HA-4SA
N2-4SA
Tumor Volume (mm3)
P = 0.4602

**Fig. S3 | TAZ identified as a major downstream effector of nSMase2. a,** Volcano plot of genechip microarray of MDA-MB-231 xenografted tumors (n = 4). **b,** RT-qPCR validation of YAP/TAZ related genes identified in microarray in MDA-MB-231 tumors (n = 4). **c,** PARADIGM integrated pathway analysis of cohort of breast cancer patients (TCGA) stratified by (quartile) nSMase2 expression. **d,** Schematic of YAP-5SA and TAZ-4SA constitutively active constructs. **e,** Validation of MDA-231 cells overexpressing YAP or TAZ via immunoblot utilizing anti-FLAG, and anti-YAP or anti-TAZ antibodies, normalized to anti-actin as a loading control. **f,** Validation of MDA-231 cells overexpressing YAP or TAZ via immunofluorescent microscopy utilizing anti-FLAG, and anti-YAP or anti-TAZ antibodies, co-stained with DAPI. **g-h,** Assessment of **g,** YAP and **h,** TAZ localization of YAP-5SA and TAZ-4SA or EV tumors via immunofluorescent microscopy with anti-YAP and anti-TAZ antibodies with DAPI used as nuclear co-stain. Quantification of nuclear localized YAP/TAZ was performed manually via blinded observer. **i, j** MDA-MB-231 cells (n = 6) overexpressing **f,** YAP1 or **g,** TAZ orthotopically implanted into the mammary fat pad of NSG mice. Tumor volume (in 2 dimensions) was measured with calipers over duration of the experiment. Tumor volume (in 3 dimensions) was measured with calipers at endpoint. **b,g-j** Data are representative of mean ± SEM of 3 or 4 independent biological replicates unless otherwise specified. *P* values obtained via student's *t*-test or two-way ANOVA + Tukey multiple comparisons test.

80 **FIGURE S4**

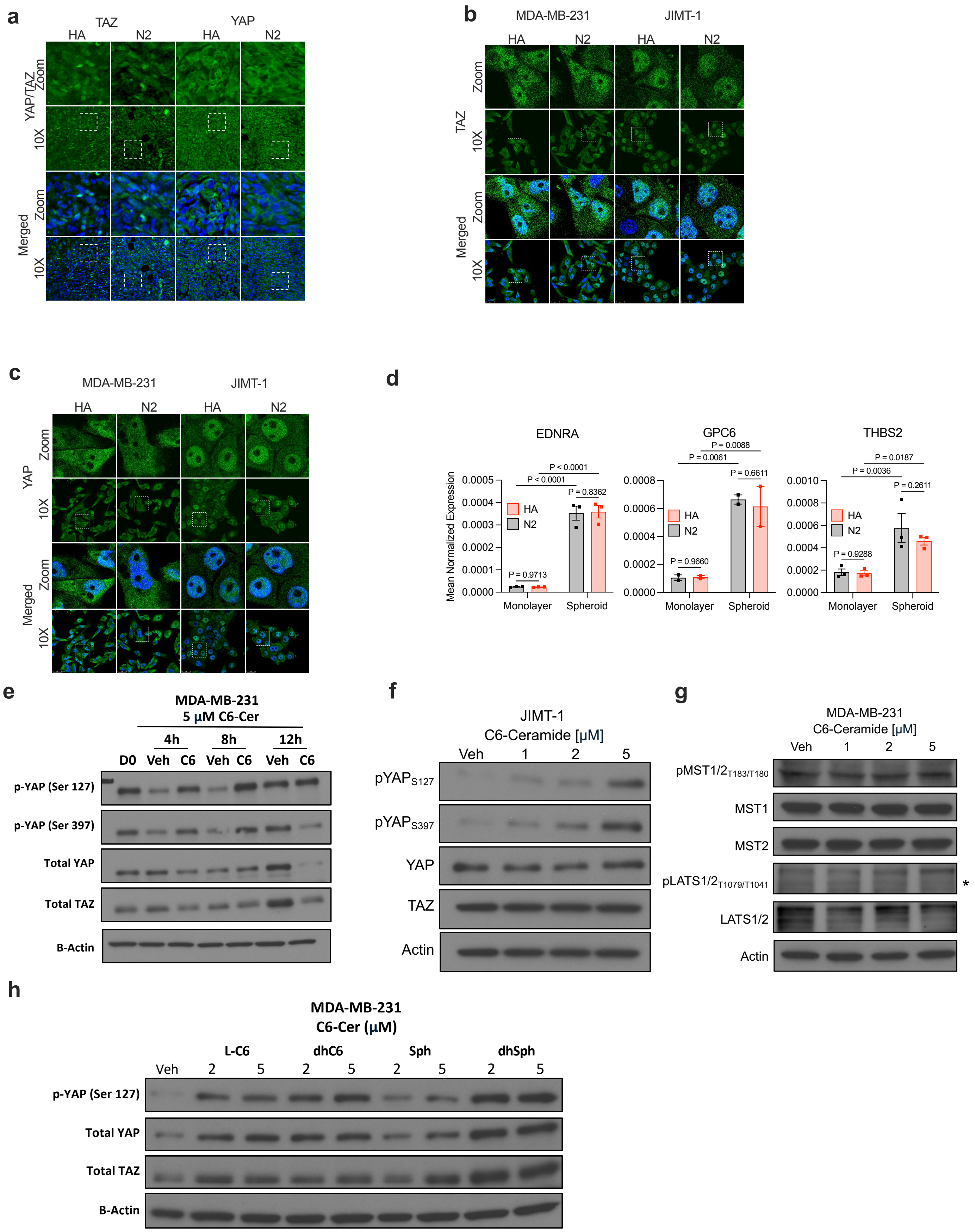


81

82

**Fig. S4 | nSMase2–Cer axis suppresses TAZ nuclear localization. a-b,** Assessment of YAP/TAZ localization via immunofluorescent microscopy with anti-YAP and anti-TAZ antibodies with DAPI used as nuclear co-stain. Quantification of nuclear localized YAP/TAZ was performed manually via blinded observer. **a,** MDA-MB-231 tumors harvested at early time point (17 d post-inoculation). **b,** MDA-MB-231 and JIMT-1 cells in monolayer. **c,** MDA-MB-231 cells in monolayer and suspension culture assessed for downstream YAP/TAZ targets via RT-qPCR. **d-e,** Immunoblot of MDA-MB-231 or JIMT-1 cells treated with C6-Cer at various doses and time points normalized to actin. **f,** Immunoblot of MDA-MB-231 cells treated with SL analogues at various doses (8 h) normalized to actin. **a-c** Data is representative of mean ± SEM of 3 or 4 independent biological replicates unless otherwise specified. *P* values obtained via student's *t*-test or two-way ANOVA + Tukey multiple comparisons test.

**FIGURE S5**

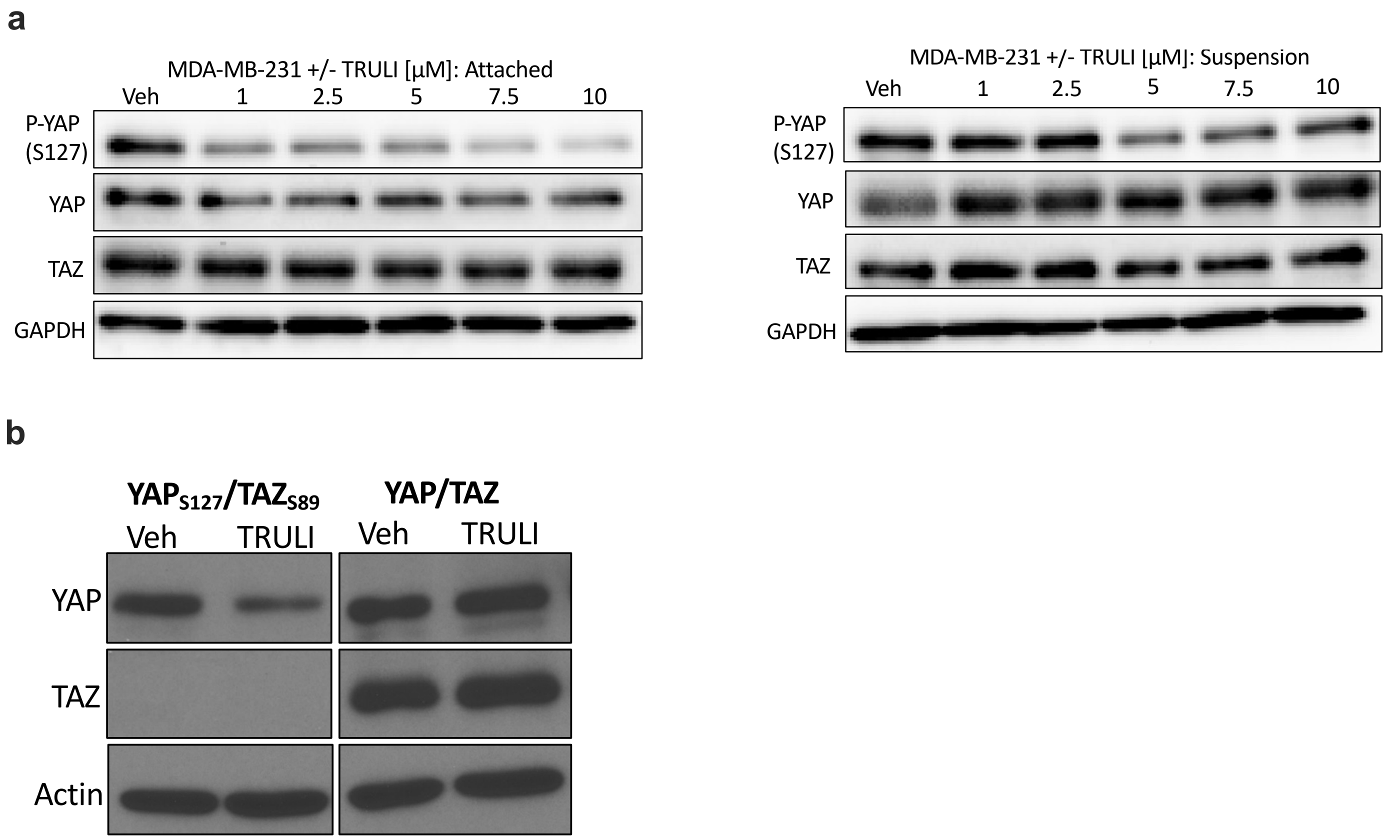


**Fig. S5 | Tumor suppressive effects of nSMase2 require LATS kinase activity. a,** Immunoblot of MDA-MB-231 cells in monolayer or suspension, treated with TRULI at various doses (24 h) normalized to GAPDH. **b,** Immunoblot of MDA-MB-231 cells 3D culture, treated with TRULI (5 µM, 48 h) normalized to actin. **a,b** Data are representative of mean ± SEM of 3 independent biological replicates unless otherwise specified.
